# Using *Drosophila* behavioral assays to characterize terebrid venom-peptide bioactivity

**DOI:** 10.1101/391177

**Authors:** Anders Eriksson, Prachi Anand, Juliette Gorson, Corina Grijuc, Elina Hadelia, James C Stewart, Mandё Holford, Adam Claridge-Chang

**Affiliations:** Institute of Molecular and Cell Biology, 61 Biopolis Drive, Singapore 138673; Department of Chemistry, Hunter College Belfer Research Center, 413 E 69^th^ St New York, NY 10021; Department of Biochemistry, Weill Cornell Medical College, Cornell University, New York, NY 10021; Division of Invertebrate Zoology, The American Museum of Natural History, New York, NY 10024; Program in Biology, The Graduate Center, City University of New York, New York, NY 10016; Program in Chemistry & Biochemistry, The Graduate Center, City University of New York, New York, NY 10016; Duke-NUS Medical School, 61 Biopolis Drive, Singapore 138673; Department of Physiology, NUS Yong Loo Lin School of Medicine, Singapore 138673

## Abstract

The number of newly discovered peptides from the transcriptomes and proteomes of animal venom arsenals is rapidly increasing, resulting in an abundance of uncharacterized peptides. There is a pressing need for a systematic, cost effective, and scalable approach to identify physiological effects of venom peptides. To address this discovery-to-function gap, we developed a sequence driven:activity-based hybrid approach for screening venom peptides that is amenable to large-venom peptide libraries with minimal amounts of peptide. Using this approach, we characterized the physiological and behavioral phenotypes of two peptides from the venom of predatory terebrid marine snails, teretoxins Tv1 from *Terebra variegata* and Tsu1.1 from *Terebra subulata*. Our results indicate that Tv1 and Tsu1.1 have distinct bioactivity. Tv1 (100 μM) had an antinociceptive effect in adult *Drosophila* using a thermal nociception assay to measure heat avoidance. Alternatively, Tsu1.1 (100 μM) increased food intake. These findings describe the first functional bioactivity of terebrid venom peptides in relation to pain and diet and indicate that Tv1 and Tsu1.1 may, respectively, act as antinociceptive and orexigenic agents. Tv1 and Tsu1.1 are distinct from previously identified venom peptides, expanding the toolkit of peptides that can potentially be used to investigate the physiological mechanisms of pain and diet.

## Introduction

Venomous animals use their expansive venom arsenal to disrupt the physiology of other animals for both defensive and predatory purposes. Due to the energetic cost of venom and the need for a fast-acting biological effect, venoms have evolved into cocktails of molecules with potent neurotoxic, haemophilic, and cytotoxic activities ^1^. Specifically, venoms contain numerous, diverse peptides, many with highly specific bioactive properties that have proven useful as pharmacological therapeutics ^2–4^. Currently, six venom-derived peptides are commercially available drugs: ziconotide for pain^5^; exenatide for diabetes ^6^; bivalirudin for anticoagulation ^7^; captopril for hypertension ^8^; and eptifibatide and tirofiban for coronary syndrome ^9^. Venom-peptide research and drug discovery has increased exponentially with the advance of genomic-transcriptomic sequencing and proteomic mass-spectrometry ^10^. However, translating newly discovered venom peptides into commercial therapeutics is challenging ^11^. A variety of high-throughput electrophysiology, fluorescence, and radioactivity-based bioassays have been used to accelerate characterization of novel venom peptides ^12^. Lacking from these is the ability to efficiently characterize the physiological properties of the enormous diversity of venom peptides being identified. There is a need for new whole-animal *in vivo* methods that are applicable to scant quantities of venom peptides.

A striking example of the discovery-to-function gap in venom peptides is the identification and characterization of peptides from marine snail venom. Predatory marine snails of the conoidean family, which includes cone snails (Conidae), terebrids (Terebridae), and turrids (Turridae), use venom peptides to prey on fish, worms, and other mollusks ^6,11,13^. Cone snail venom peptides molecularly target ion channels and receptors, and in the case of cone snails, have shown remarkable selectivity for certain channels and receptors ^14–16^. Additionally, the compact size of conoidean venom peptides and their enhanced disulfide-bridged stability make them good candidates for pharmaceutical drug discovery and development ^17,18^. There are over 15,000 species of conoideans, with an estimated venom-peptide reservoir of over a million compounds ^19^. However, less than 2% of conoidean venom peptides have been functionally characterized to date. Originally identified with assays in mice, Prialt (ziconotide) is the first conoidean venom drug, and it is used to treat chronic pain in HIV and cancer patients ^5,20^. Ziconotide is chemically identical to the naturally occurring *Conus magus* peptide (MVIIA), and has illuminated a new molecular target for treating pain, namely N-type calcium channels ^21,22^. Ziconotide’s success has peaked the interest of pharmaceutical companies and there are several other cone snail venom peptides, conotoxins, in various stages of pharmaceutical drug development ^5,23–27^. Given this, devising new cost-effective strategies to identify the bioactivities of snail-venom peptides could advance their use as biomedical physiological tools and development as drug candidates.

While conotoxins have received a lot of attention, the venom peptides from terebrid snails are less studied, representing an untapped resource. Terebrid transcriptomes and proteomic data reveal that similar to conotoxins, the venom peptides from terebrid snails—teretoxins—are highly structured and disulfide-rich; two features that benefit stability and pharmacokinetics ^28–30^. However, it is important to note that while evolutionary similar, teretoxin and conotoxin differ in size, structural integrity and complexity, suggesting that they might be applicable to different molecular applications ^29^. The potential of novel attributes of teretoxins for combating human disease and disorder demands efficient strategies to characterize these promising bioactive peptides ^17,31^.

Pain and obesity are two major therapeutic areas for which an improved comprehensive pipeline that includes high-throughput physiological screening of teretoxin venom peptides could have an impact. In the U.S., the over-prescription of opioid pain medications has led to an epidemic of addiction and overdose for which treatment remains elusive ^32,33^. Remarkably, a potential solution to non-addictive pain therapies is the snail venom drug ziconotide. As ziconotide’s molecular site of action is N-type calcium channels and not opioid receptors, it is not addictive ^34^. However, ziconotide does not cross the blood brain barrier and has to be delivered to the central nervous system via intrathecal injection, an invasive delivery method that limits its use ^35,36^. Even with its limitations, ziconotide is a paradigm shifting breakthrough that demonstrates there is an alternative non-addictive strategy for treating pain. As a result, here we examined teretoxins to identify new snail venom peptides that may have nociceptive properties and are peripherally active.

Similar to the upward trend in pain treatment, in the past 30 years the prevalence of obesity has reached epidemic proportions where the morbid obesity rates continue to rise ^37^. Obesity is recognized as a major public health concern and is associated with numerous complications such as diabetes, cardiovascular diseases and various forms of cancer ^38–40^. Obesity is a complex disorder caused by an imbalance between energy intake and the expenditure and the interaction between predisposing genetic and environmental factors ^41–43^. Several peptidergic systems within the central nervous system and the periphery are known to regulate energy homeostasis. However, the specific gene-gene and gene-environment interactions involved in this process remain unclear. A proposed treatment to combat obesity is being able to restore the energy homeostasis as a result of a dysregulation in the peptidergic system ^44^. However, despite several efforts, the cause of obesity still remains inconclusive and the only effective treatment against obesity currently is gastric bypass surgery ^45^. New compounds that can be used to investigate the cellular physiology of pain and obesity are needed in order to develop effective treatment.

In the search for new bioactive compounds that may be effective against pain and diet, we developed a physiological approach using vinegar flies (*Drosophila melanogaster*) to screen biologically active teretoxin peptides. Flies are abundant, affordable, and amenable to automated behavioral testing, making them suitable for testing venom-peptide bioactivity. *Drosophila* has been successfully used for drug screening in a number of examples: multiple endocrine neoplasia type 2 ^46^, fragile X syndrome ^47^, epithelial malignant growth ^48^, intestinal stem cellderived tumors ^49^, combinatorial therapy ^50^, and life span ^51^. We focused on fly thermosensation and feeding, behaviors that have been used to model pain and obesity. Our approach uses *in vivo* pain and diet fly assays to characterize venom peptides from terebrid snails (Figure 1). The strategy requires minimal amounts of venom peptide, by using a genetically tractable small-animal model. To establish the viability of this approach, we used several fly assays to characterize two teretoxins: Tv1 from *Terebra variegata* and Tsu1.1 from *Terebra subulata*. The two peptides had distinct properties: Tv1 displayed reduced avoidance towards noxious heat; Tsu1.1 increased the number of feeding bouts. The disparate bioactivity of Tv1 and Tsu1.1 supports the robust nature of our systematic fly behavioral assays for identifying new venom peptides that may have therapeutic relevance.

**Figure 1.**
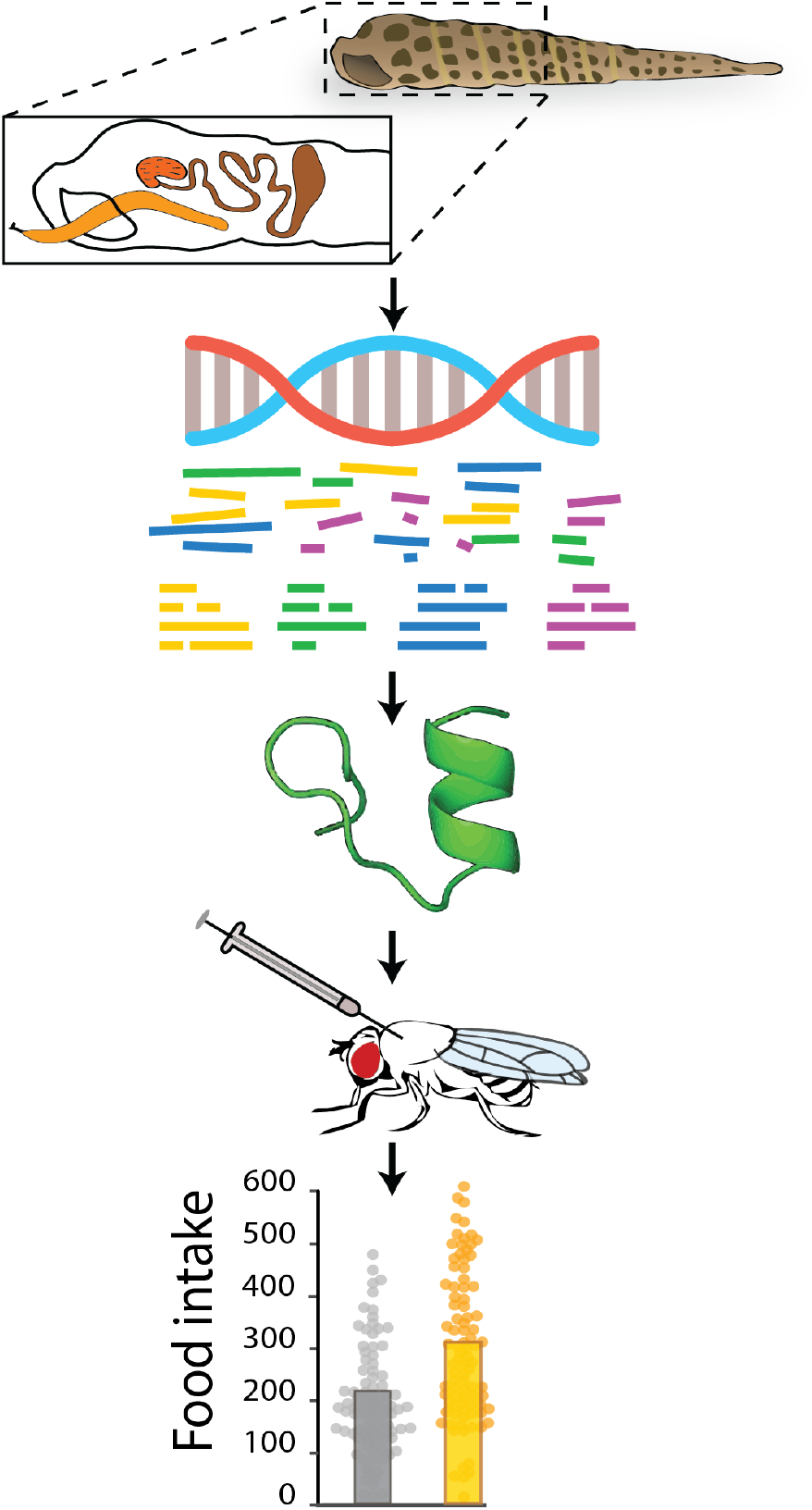
Schematic representation showing the pipeline used in this study. Venom glands are dissected from terebrids in the field. RNA is extracted in the lab and sequenced using next-generation sequencing. Transcriptomic data are assembled and searched for peptide candidates. Selected peptides are chemically synthesized, oxidized and injected into flies. Bioactivity is measured in several behavioral assays.

## Results

### Tsu1.1 is a novel teretoxin peptide

We used a bioinformatic pipeline to sequence, assemble and search the *Terebra subulata* venom-gland transcriptome for novel-peptide candidates (Figure 2A). One of these candidates was a sequence with 21 amino acid residues (SAVEECCENPVCKHTSGCPTT), which we called *Terebra subulata* teretoxin 1.1 (Tsu1.1) (Figure 2B). We compared Tsu1.1 with another peptide, *Terebra variegata* teretoxin 1 (Tv1) which was previously characerized in the Holford group^54^. Teretoxins Tv1 and Tsu1.1 have distinct cysteine frameworks—a feature that is used to group conotoxins into functional groups— which suggests they might have different molecular targets ^31,55^. Tv1 is arranged CC-C-C-CC (Framework III) while Tsu1.1 is arranged CC-C-C (Framework I) (Figure 2C). In cone snails, Framework III conotoxins are in the M superfamily, a group of peptides that target either sodium (Na+) or potassium (K+) channels ^56^. By contrast, Framework I conotoxins are in the A superfamily, which target either adrenergic receptors or nicotinic acetylcholine receptors (nAChR) ^56^. An alignment of Tv1 and Tsu1.1 with known conotoxins confirms that, beyond their cysteine scaffold, and a conserved proline in Tsu1.1, Tv1 and Tsu1.1 are not homologous to members of their respective conotoxin superfamilies and are distinct from known conotoxins (Figure 2C) ^54^.

**Figure 2.**
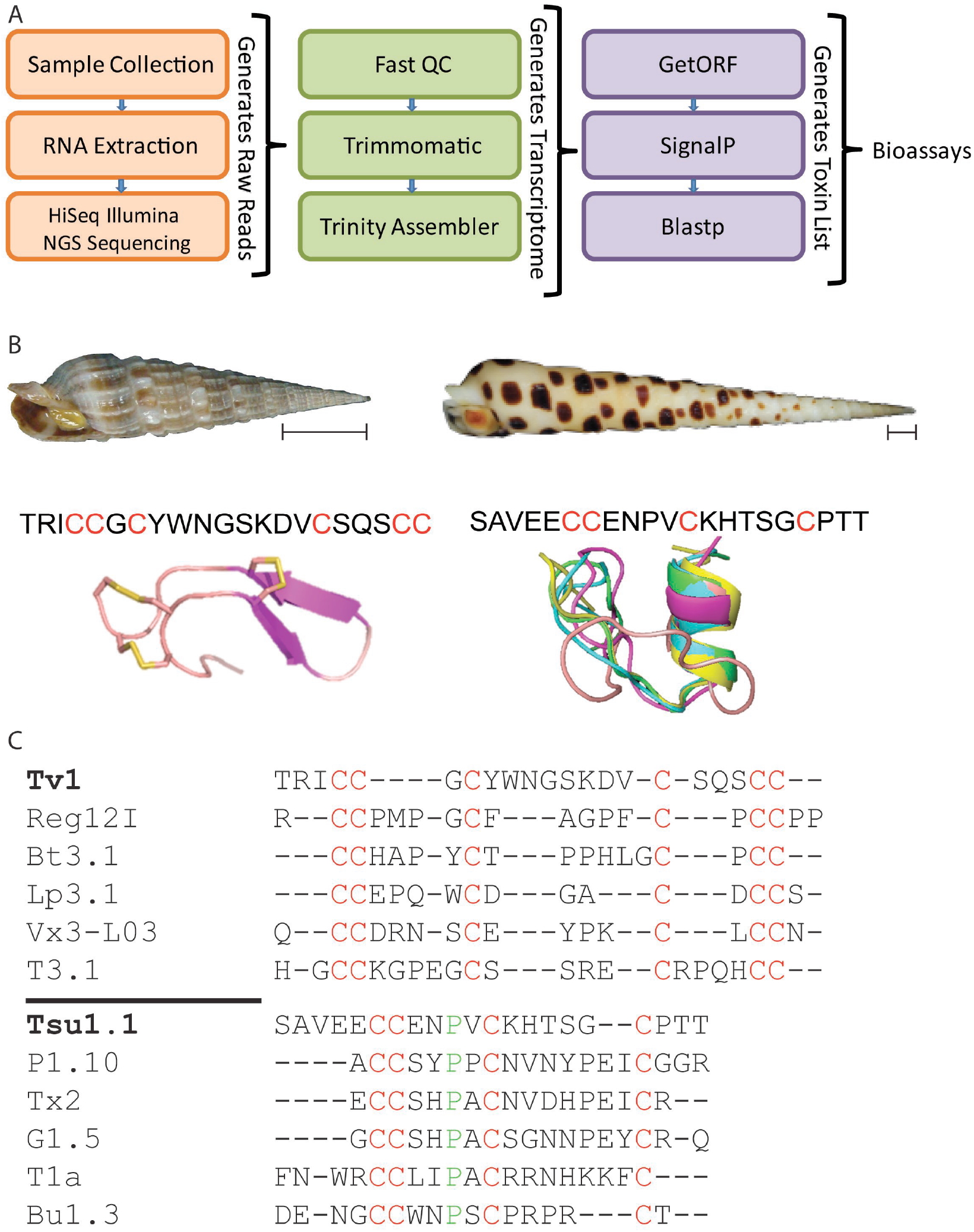
Comparison of Tv1 and Tsu1.1. A. The bioinformatic pipeline used to identify Tsu1.1. The orange box indicates the initial steps to generate reads through next-generation sequencing (NGS), the green shows the process to assemble the terebrid venom gland transcriptome, and the purple represents the final steps that lead to the discovery of teretoxins. B. Shell images of *Terebra variegata* (left) and *Terebra subulata* (right), scale bar represents ~1 cm, followed by the sequence of each venom peptide, with cysteines highlighted in red, followed by the NMR structure of Tv1^46^ and the predicted I-TASSER structure of Tsu1.1. C. An alignment of Tv1 and conotoxins in the M-superfamily, followed by an alignment of Tsu1.1 and conotoxins in the A-superfamily. Green represents shared amino acids, while red represents shared cysteine framework.

### Tv1 and Tsu1.1 have minimal effects on mortality

We comparatively examined the bioactivity of Tsu1.1 and Tv1 teretoxin peptides. In order to investigate the physiological activity of the two peptides, we determined the levels at which the two peptides had lethal toxicity. Initial concentration was decided based on previous studies investigating the bioactivity of Tv1 in polychaete worms, where it was found that Tv1 caused a partial paralysis at a concentration of 20 μM^46^. However, an initial test with injections of 100 μM peptide (the highest dosage tested) found that—even at this concentration—neither peptide had major mortality effects in flies (Figure S1A–B). Based on these results, we used injections of either 100 μM or 20 μM in subsequent experiments to explore Tv1 and Tsu1.1’s non-lethal bioactivities.

### Tsu1.1 injections induce mild hypoactivity

While Tv1 had only trivial effects on activity, Tsu1.1 injections had noticeable, though small, effects on multi-day daytime and nighttime activity (Figure 3A, C). Over a five-day assay, Tsu1.1-injected flies walked 27% less (−226.93 mm [95CI - 472.1, 29.29], *g* = −0.34, *P* = 0.04) than controls during the daytime (Figure 3B), or a Δdistance of −226.93 mm ([95CI −472.1, 29.29], *g* = −0.34, *P* = 0.04) (Figure 3B). We investigated if the Tsu1.1 effect was associated with functional motor deficits using a climbing assay ^57^. Upon injection with Tsu1.1, the flies exhibited only small or negligible differences in climbing index (Figure 3C and 3D); this result suggests that Tsu1.1 does not diminish motor coordination.

**Figure 3.**
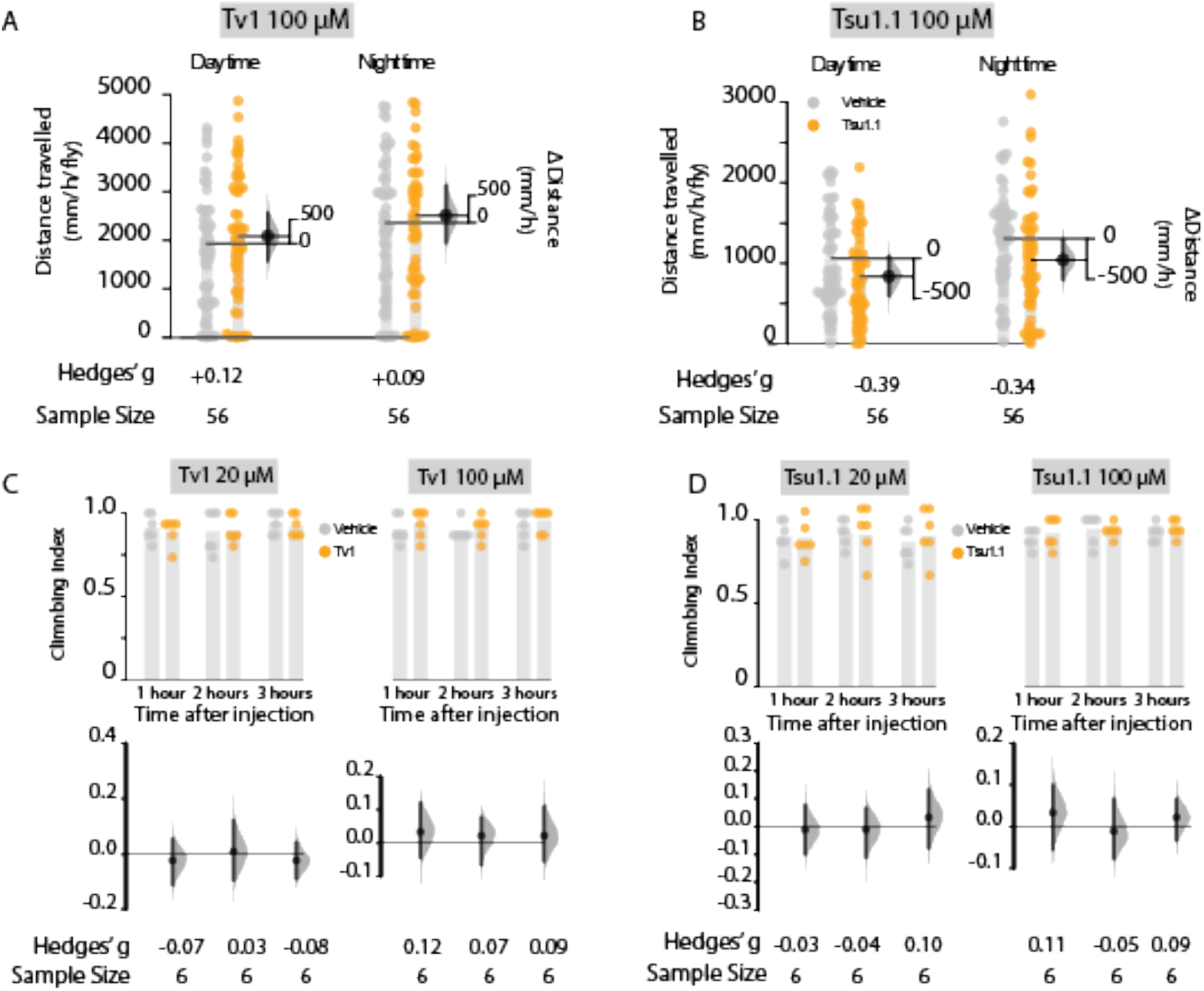
Effects of Tv1 and Tsu1.1 on activity and motor coordination of male Drosophila. Activity was recorded for 5 consecutive days using automated video tracking and measured as the distance travelled per hour (mm/h/fly). Controls were maintained during the same conditions as the teretoxin-injected flies with the only difference that they were injected with PBS. **A.** Activity assay of Tsu1.1; **B.** Activity assay of Tv1; **C.** Climbing index measured the short term effects of the toxins as well as possible disruptions in their motor coordination. The climbing assays of Tsu1.1 revealed only trivial effects on climbing ability. **D.** Climbing assay of Tv1, all at 20 μM and 100 μM. All error bars represent 95% CI. The numbers indicated below each bar denote the standardized effect size (*g*). The climbing index was done with at least 15–20 male flies in each vial. Control data are depicted with grey dots, and injected-fly data with orange dots.

### Injection of Tv1 reduces heat avoidance

Teretoxin Tv1 has the same cysteine framework as known M superfamily conotoxins, members which have been characterized for targeting voltage-gated sodium channel (VGSC) subtypes Na_v_1.8 and Na_v_ 1.9. These channels have been associated with controlling neuropathic pain ^58–60^, and some M-superfamily peptides are potential antinociceptives ^14,16^. Based on its cysteine-framework similarity to M-superfamily conotoxins, we examined if Tv1 had antinociceptive activity. We used an assay that measures *Drosophila* heat avoidance, that has been applied as a model of acute pain^61^. The assay monitors innate avoidance of noxious (46°C) surfaces (Figure 4A). The heat-avoidance assay was performed on flies injected with 100 μM of Tv1 and Tsu1.1, and analyzed at 1 and 2h time intervals after injection). One hour after injection, control flies injected with PBS had a heat avoidance of 70.9 [95CI 65.4, 76.4]; Tv1-injected flies exhibited a markedly reduced avoidance response of 58.6 [95CI 53.0, 64.2]. This represents a 12.33% reduction in avoidance ([95CI −20.31, −4.39], *g* = −1.02, *p* = 0.005) after one hour; in terms of standardized effect sizes, this is a large effect ^62^. By the second hour, this response diminished to −4.97% ([95CI −14.34, 4.7], *g* = −0.43, *p* = 0.33) (Figure 4B). For Tsu1.1, injected flies displayed heat avoidance that was similar to controls: 70.5% ([95CI 65.5, 75.5]), or an avoidance change of −2.3% ([95CI −10.4, 5.8], *g* = −0.26, *p* = 0.58v). These findings indicate that Tv1 reduces fly sensitivity to noxious heat.

**Figure 4.**
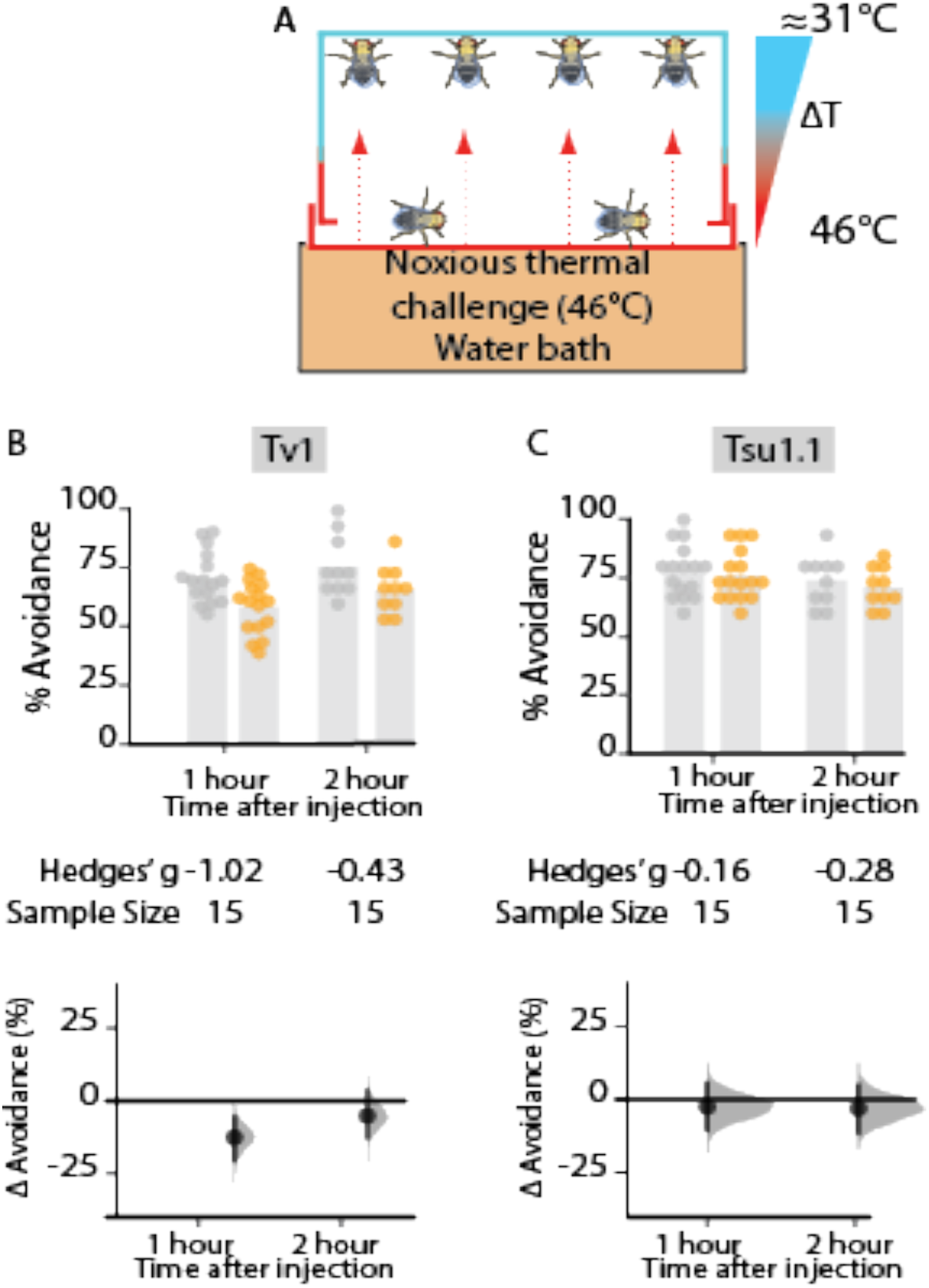
Thermal nociception in adult *Drosophila* is diminished by Tv1. A thermal heat assay used to measure the antinociceptive effect of Tsu1.1 and Tv1 on adult flies. A. Schematic representation of the assay measuring thermal nociception. The assay measures the avoidance from a noxious (46°C) surface. B. Injection with the higher concentration of Tv1 produced a large reduction in heat avoidance in the first hour: Δ = - 12.33% [95CI −20.31, −4.39], *g* = −1.02, *p* = 0.005, N=15. Hedges’ *g* is a standardized effect size. C. Avoidance behavior in flies injected with two different concentrations of Tsu1.1: the control flies and the experimental flies had a similar heat avoidance. Control data are depicted in grey dots, and injected-fly data in orange.

### Tsu1.1 administration increases food intake

Feeding and foraging behaviors strongly rely on a neuronal and endocrinological connectivity through the gut-brain axis that also includes a peripheral regulation ^63–65^. Thus, while we do not know whether teretoxins Tv1 and Tsu1.1 can cross the fly blood-brain barrier, the injected peptides could affect feeding via peripheral systems. Consequently, we assayed feedings over a 6 h time period using capillary-feeder assay^66^ that was adapted for video tracking of single-fly feeding. In the Tv1 experiment, almost no difference in food intake was seen after injection: food intake decreased only −6.13 nl ([95 CI −23.10, 35.08 ], *g* = - 0.07, *P* = 0.7) (Figure 5E). In the Tsu1.1 experiment, while control flies injected with PBS drank an average of 214.0 nl [95 CI 180.12, 247,88] the Tsu1.1 injected flies drank 307.4 nl [95 CI 270, 344.8]: 46% more liquid: an increase of +93.34 nl [95 CI 41.61, 141.485 ], *g* = +0.55, *P* = 0.0002 (Figure 5A).

**Figure 5.**
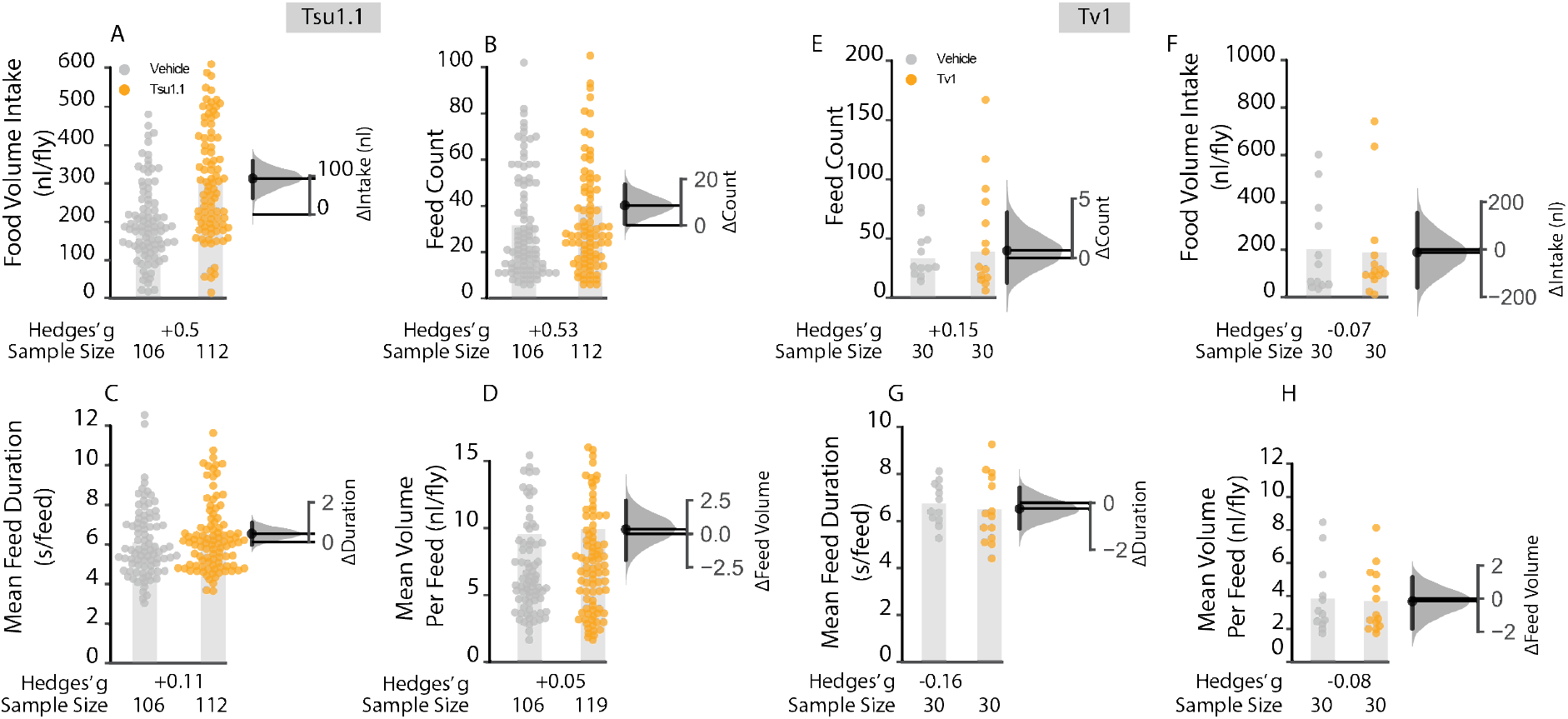
Tsu1.1 has effects on food intake by increasing meal frequency. A single-fly feeding assay was used to assess total food intake, the average number of meals (meal duration, and meal size (over a 6 h period in 5-7 day old adult Canton-S males. **A.** Flies injected with Tsu1.1 toxin consumed more: + 93.3 nl [95 CI 41.6, 141.49 ], g = +0.55, P = 0.0002, N = 106, 112. **B.** Average number of feed bounts where Tsu 1.1 injected flies displayed a dramatic increase: Δfeeding bouts +8.64 [95 CI 0.53, 17.23], g = +0.53, p = 0.047, N = 106, 112. **C-D.** Mean meal duration and volume per feed showed only minor alterations after injection of Tsu1.1. **E-H.** The feeding behavioral effect of Tv1 dissected into volume intake, feed count, feed duration and volume per feed. All metrics showed only trivial differences. The data represent the mean differences with their 95% CI. Control data are depicted in grey, and injected-fly data in orange. The numbers below each column denote the effect size and sample sizes (N) for each experiment.

### Tsu1.1 increases meal frequency

To determine the behavior that produces the increased consumption, we examined the aspects of foraging behaviour (Figure 5B). Consummatory behaviors in individual flies were analyzed over a 6 h period, allowing us to determine three additional feeding parameters: number of meals, the average meal size, and the meal duration (Figure 5B–D). Injection of Tv1 had no effect on any of the parameters analyzed (Figure 5E–H). However, Tsu1.1 injection elicited a increase in meal count: while control flies had 31.56 meals per fly [95 CI 27.16, 35.96], flies injected with Tsu1.1 consumed 40.20 meals [95 CI 36.64, 43.76]. This was a +8.64 meals ([95 CI 0.53, 17.23], *g* = +0.53, *p* = 0.047) or an increase of 27%. Feed volume and the total feed duration was also measured, both of which showed very small increases (Figure 5C–D). Thus, the increased feed volume appears to arise largely as a result of an increased number of meals, with very minor contributions from increased meal duration and volume.

## Discussion

This work demonstrates the potential of *Drosophila* behavioral assays to be used to analyze the physiological bioactivity of venom peptides. Specifically, we characterized the physiological effects of two teretoxins, Tv1 and Tsu1.1 from *Terebra variegata* and *Terebra subulata* respectively on thermosensation and feeding, behaviors that have been used to model pain and obesity. Pain and diet-related disorders are two areas for which new bioactive compounds are needed to advance therapeutic development.

Previously, Tv1 injected into *Nereis virens*, a polychaete worm, has been shown to induce partial paralysis ^54^. This effect was absent in *Drosophila* (Figure 3A and C), suggesting that the Tv1 binds to targets which are not conserved between polychaetes and insect. However, our findings did show that injected Tv1 reduces fly heat avoidance (Figure 4). This is the first demonstration that teretoxins can have an antinociceptive effect. A possible mechanism of action for Tv1-induced antinociception in flies is by modulation of TRP (Transient Receptor Potential) channels. TRP channels are like physiological canaries in the mine, they sense external stimuli and and serve as an early warning system for the organism. TRP channels are found throughout animal species including humans, mice, worms and zebrafish (Figure 6). It has been previously shown that *Drosophila* TrpA1 is a thermoreceptor and a critical mediator of nociception in flies and mammals ^67,68^. Prior reports suggest *Drosophila* TRP is required for preserving light response, whereas external noxious heat stimuli (>40^0^ C) activates TrpA1, Pyrexia, and Painless channels ^69–72^. The decreased sensitivity to noxious heat observed in our heat avoidance assay for Tv1 treated flies suggests it is possible that Tv1 inhibits one of the three heat-activated TRP channels. While the Tv1-induced decrease was small, it is a significant finding in that Tv1 was peripherally administered, therefore Tv1 could serve as a scaffold for designing novel peripherally active antinociceptive peptide compounds.

**Figure 6.**
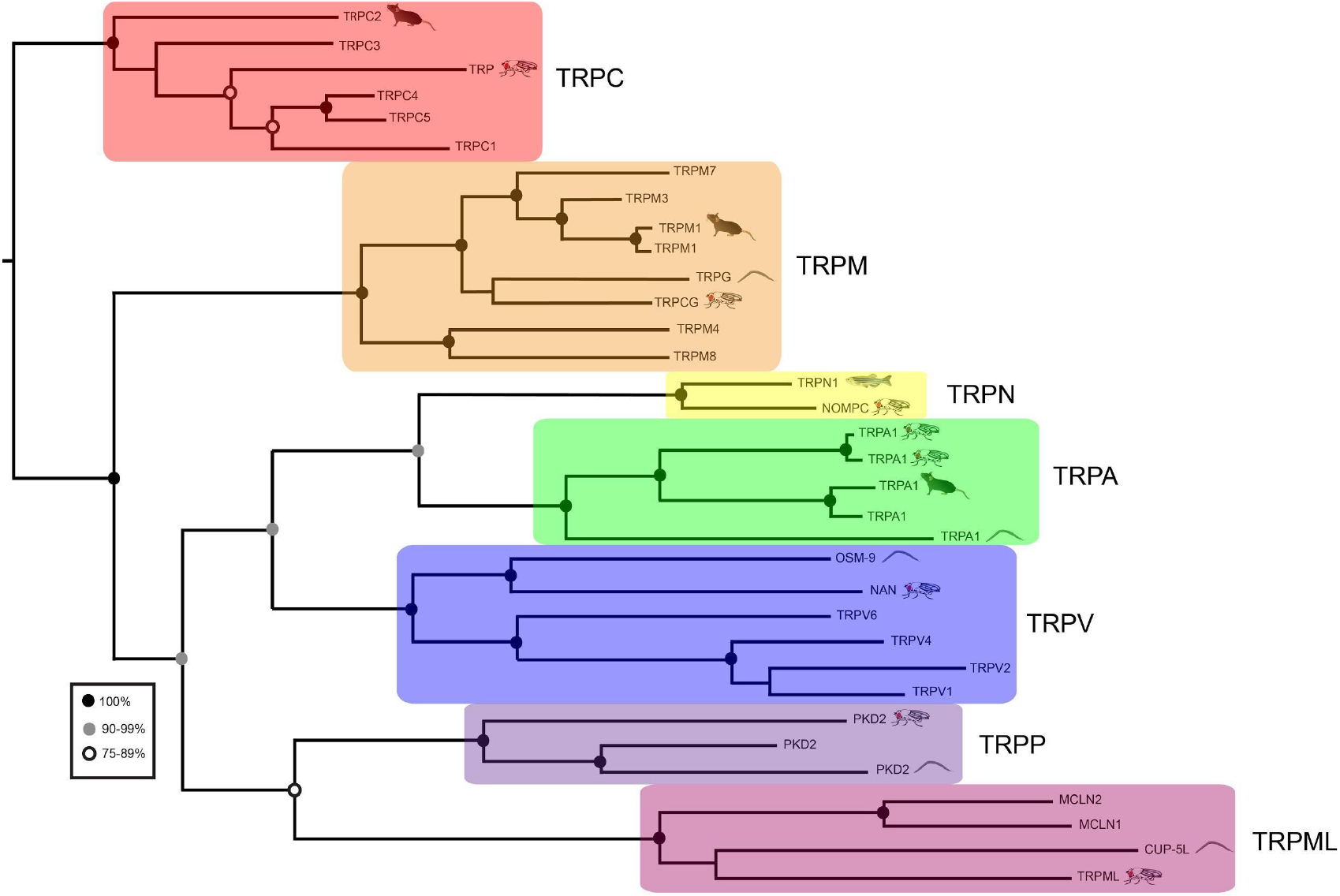
Comparison of TRP channels across the tree of life. Maximum-likelihood phylogeny showing the relationship between members of the human TRP-channel superfamily and members of *Caenorhabditis elegans* (worm), *Drosophila melanogaster* (fly), *Danio rerio* (zebrafish) and *Mus musculus* (mouse) TRP channels. *Drosophila* TRP channels associated with pain, such as TrpA1, are also found in humans, mice, and worms.

Alternative to Tv1, Tsu1.1 has an orexigenic effect, increasing food consumption by 46%. Using a single-fly feeding assay with automated food-consumption tracking allowed us to dissect several aspects of feeding: intake, meal count, meal volume and meal duration. Of these, increased meal count appeared to account for some of the food intake increase, while meal volume and duration underwent only small changes. This increase in feed count does not account for the entire increase in the total consumption. As Tsu1.1 was administered by injection, it is possible Tsu1.1 affects the peripheral system of the fly and regulates feeding via a neuroendocrinological pathway. Several studies have been tried to isolate the effect of neuromodulators on feeding ^73,74^. A majority of these neuromodulators have a negative effect on feeding most likely due to disruptions of its motor coordination. However, previous studies indicate activation of serotonin, octopamine and dopamine causes an increase in feeding ^73,75–77^. Further analyses are needed to determine which, if any, of these systems are affected by Tsu1.1.

## Conclusions

We developed a sequence driven, activity-based hybrid approach using venom transcriptomic analysis and behavioral characterization in *Drosophila melanogaster* to identify venom-peptide bioactivity. Application of our method found that Tv1 reduces heat avoidance in fly, suggesting that it has a antinociceptive effect, and also indicated that Tsu1.1, a novel teretoxin, modulates feeding behavior. The orexigenic effect of Tsu1.1 arose primarily from increased meal frequency, with additional effects from meal duration and volume. Taken together, our findings illustrate the utility of *Drosophila* assays to identify the bioactivity of novel venomics-derived peptides. This phenotypic-screening approach can help to realize the enormous potential that terebrid snail venom peptides have as compounds for characterizing cellular physiology, and as candidate therapeutics.

## Methods and materials

### Identification of venom peptides

Tsu1.1 was discovered using a transcriptomics and bioinformatics pipeline (Figure 2A). RNA was extracted from the venom duct of *Terebra subulata* using an RNeasy Micro kit (Qiagen). RNA was sequenced with an Illumina HiSeq 2500 with a multiplexed sample run in a single lane, using paired end clustering and 101 × 2 cycle sequencing. Raw reads were first processed on Fast QC to assess the quality, looking for a phred score ≥30. Low quality reads and adapters were trimmed using Trimmomatic. The *de novo* assembly program Trinity ^78,79^ was used with default parameters to assemble the transcriptome. Once a transcriptome was assembled, EMBOSS’ GetORF ^80^ was used to translate transcripts into Open Reading Frames (ORFs). SignalP ^81^ was used to predict signal sequences in the translated ORFs. As teretoxins are secretory peptides, they are all associated with a signal sequence that is cleaved off from the mature toxin sequence before envenomation. Only those ORFs with signal sequences were analyzed using the basic local algorithm search tool (BLAST) ^82^ and an in-house toxin database to identify putative toxins homologous to already known toxins. The sequence of Tsu1.1 was found to be SAVEECCENPVCKHTSGCPTT.

### Synthesis and oxidative folding of Tv1

As teretoxins are present only in miniscule quantities in the venom, a greater quantity of the linear peptide Tv1 was chemically synthesized and purified by previously described methods and further oxidatively folded into biologically active form ^54^.

### Synthesis and purification of Tsu1.1

Tsu1.1 peptide with a sequence of SAVEECCENPVCKHTSGCPTT was synthesized by microwave assisted Fmoc solid-phase peptide synthesis on a CEM Liberty synthesizer using standard side chain protection. Following treatment of peptidyl resin with Reagent K [92.5% TFA (Trifluoroacetic acid), 2.5% TIS (Triisopropylsilane), 2.5% EDT (1,2 Ethanedithiol) and 2.5% water, 4h)] and methyl tertiary butyl ether (MTBE) precipitation, crude Tsu1.1 checked for purity on Agilent UHPLC machine and eluted using a linear gradient from 0% to 55% buffer B (80% Acetonitrile with 20% water) in 3.5 min. The identity of synthesized peptide was confirmed by molecular mass measurement of purified peptide using 6520 Agilent Q-TOF LC-MS (Figure S2 and S3).

### Oxidative folding of peptides

A one-step thiol-assisted oxidation was used to prepare folded Tsu1.1 peptide. The linear peptide (20 μM) was incubated in 0.1 M Tris -HCl, 0.1M, NaCl, 100 μM EDTA,1 mM GSH (reduced glutathione), 1 mM GSSG (oxidized glutathione), pH 7.5. The folding reaction was terminated by acidification with 8% formic acid at 15 min, 30 min, 1, 2, 3, 4, and 24 h and the folding yield monitored using UHPLC. A preparative scale folding reaction was then conducted at an optimized time of 1 h, and the folded peptide was purified using X-Bridge semi-preparative column (waters). Elution was carried out at 5 ml/min with 0% buffer B and 100% buffer A for the first 5 min, then increasing buffer B to 35% in 35 min. The purity was confirmed using Agilent poroshell UHPLC column and the molecular mass of the oxidized peptide was confirmed by LC-MS. The oxidatively folded peptide was used in all the experiments (Figure S4 and S5).

### Structure prediction of Tsu1.1

The membrane protein topology and signal peptide prediction was made using TOPCONS according to the online instructions ^83^. Model structure was made by using the I-TASSER web application ^84,85^. The quality of the model predicted by I-TASSER is verified using two different score systems: TM-score and a root mean square deviation. These two scoring systems gives an estimate how close the model is to the native structure and both of these values are computed from a confidence score (C-Score) ^84,86^. For the TM-score a value above 0.5 usually implies a correct topology for the model ^87,88^. The top five high scoring structures based on C-score from the I-TASSER prediction were superimposed using PyMol.

### Multiple sequence alignment

The ConoPrec tool, which is available on the ConoServer website was used to identify similar sequences recorded in the ConoServer database ^89,90^. The multiple sequence alignment was performed using ClustalW software ^91^.

### Fly stocks

All *Drosophila melanogaster* flies used in the experiments were male Canton-Special aged 5–9 days at the onset of the experiment. Experimental flies were maintained at 25°C, 60–70% relative humidity, under 12:12 h light and dark cycles for 4 to 7 days before the experiment day. If starvation was required, flies were wet starved for 24 h before experiments. Wet starvation was performed by keeping flies in a plastic vial with 2% agar.

### Teretoxin peptide injection

Concentration of the toxins was determined based on previous studies investigating the bioactivity of Tv1^54^ which concluded Tv1 to cause partial paralysis in polychaete worms at a concentration of 20 μM. A low (20 μM) and high (100 μM) concentration was used for both peptides to obtain a range for bioactivity. Injection glass micropipettes (Nanoliter2000 borosilicate glass capillaries, World Precision Instruments, Sarasota, FL, U.S.A.) were pulled with a horizontal pipette puller (model P-1000, Sutter Instrument Co., Novoto, CA, USA) creating a 11–17 μm wide opening. Freshly pulled needle was blunted by pressing the tip through a paper tissue. The injection micropipette was then mounted on a microinjector (Eppendorf Femtojet Express 5248) at an injection pressure of 300 kPa. Flies were anesthetized using CO_2_ and kept sedated throughout the entire injection process (maximally for 5 minutes). Anesthesia was maintained by keeping the flies on a pad consisting of porous polyethylene. A volume of 0.2 μl was injection by a pulse pressure of 300 kPa under stereomicroscope. Typically, the injection site was made intra-thoracically beneath the supraalar bristles (SA1, SA2) and the presutural bristle, into the area between between the mesopleura and pteropleura.

### Locomotion assay

Flies were tested using an automated video tracking software measuring the distance traveled of the flies. Flies were placed in 5 mm × 65 mm glass tubes with food placed at one end of the tube and a cotton plug in the other, allowing proper respiration of the flies. The food was allowed to air dry for one hour (to prevent flies from drowning in wet bait) before the flies were added. A total of 56 flies were used for each bioactive peptide with the same number of control flies injected with the vehicle. To determine phenotypic defects in the flies upon injection of Tv1 and Tsu1.1, we examined their activity using an automated video-tracking assay. The activity assay employs video tracking to measure the total distance (mm/h/fly) over five consecutive days and excludes a 24 h habituation period. The tracking assay measured the long term and short term effects of distance travelled and circadian rhythms of the flies. The flies were tracked with computer vision code in LabVIEW using standard background subtraction and centroid methods. Behavior data was analyzed and plotted with Python using Jupyter and the scikits-bootstrap, seaborn and SciPy packages. Data was collected for five consecutive days with the first day subtracted from the assay, a habituation period. The data was then averaged by hour over the entire duration of the experiment.

### Thermosensation assay

A thermosensation pain assay for adult *Drosophila* was performed as previously described ^61^. Approximately 20 flies, 5 to 7 days old were placed in sealed behavioural chamber (Petri dish measuring 35mm x 11mm; Nunclon). Flies were allowed to acclimatize to the environment for 60 min at room temperature. Using a water bath, the bottom of the chamber was heated to 46°C. The chambers were heated for 4 min before they were removed from the water and immobilized flies were counted. The percentage avoidance was calculated by counting the number of flies that failed to avoid the noxious temperature compared to the total number of flies in the chamber. The distance from the heated bottom for each fly was not taken into consideration.

### Climbing assay

Twenty-four hours before the experiment, male flies were separated from the females and transferred to a newly prepared food vial. No more than 20 flies were kept in the same vial. After the habituation period, five flies were transferred to 50 ml serological pipette tubes (Falcon, USA) that was cut to 50 mm in length. The top and bottom of the tube were sealed with parafilm with three small holes to provide ventilation. Flies were habituated to the new environment by lying flat on the surface for one hour at 25°C. The tube was tapped against a hard surface at the beginning of the experiment to place all the flies at the bottom of the vial. The time for each fly to reach the top marked point was recorded. Any flies that could not reach the top mark within 60 seconds were marked as failure to climb. Climbing index was calculated as a range from 0 to 1 for the number of flies that managed to reach the top mark within a certain time. A failure to reach the top mark gave them a score of 0 and a success a score of 1. An average for all flies was calculated for all the trials.

### Feeding assay

Male flies were anesthetized by cooling and placed in chambers for the feeding assay, where capillaries delivered liquid food (5% sucrose, 10% red colour food dye in deionized water) to the fly. The experiment was conducted within an incubator that was maintained at 22°C^66^. The individual toxins were tested on different days; all experimental conditions were kept constant in between and during each experiment with respect to: temperature, humidity, circadian time, days after eclosion and the sex of the flies. The level of the fluid was monitored using video tracking for 6 h. Each capillary was accessible by a single fly kept in a 12mm × 12mm × 2mm chamber cut from acrylic. The food intake assays for the individual fly lines were not performed concurrently but with identical duration, start time for the experiment, age and sex of flies, temperature and starvation time. For experiments involving starvation, flies were wet starved for 24 h before the start of the experiment; during wet starvation, flies were deprived of food but not water in a vial containing 1% agarose dissolved in deionized water. Different sets of flies were used for the vehicle and toxin injections, as well as for the fed and the food-deprivation.

### Determination of effects on activity and motor coordination

Flies aged for 5–7 days after eclosion were anesthetized using CO_2_ and single flies were transferred to glass tube (5 mm x 665 mm) containing food in one end and sealed with a plastic cap with the opposing end sealed with a cotton wool plug. Standard fly food was used for both the experimental and negative control. One hour prior to the start of the experiment flies were anaesthetized and injected with the toxin. The flies were allowed to recover for one hour before being transferred to the glass tubes and inserted into the arena within a light and temperature regulated incubator. The temperature was set to 22°C with a 60–70% relative humidity, under 12:12 h light and dark cycles. When comparing experimental and control flies, they were distributed with 56 control flies and 56 experimental flies on each side of the arena. The flies were habituated for 24 h before commencing the experiment. The activity was automatically measured by tracking the path of each individual fly by means of video tracking and calculated as the numbers of crosses over the midline of the glass tube along with distance travelled. To assess sleep duration a threshold for sleep was used and is according to previously published protocols for sleep assessment in *Drosophila*: a period of inactivity lasting for at least five consecutive minutes ^92^ sleep duration was measured as the cumulative amount of sleep in a 24 h period measured in minutes. As baseline activity of control animals was susceptible to change between experiments over time, each peptide was tested in otherwise identical animals assayed in parallel.

### Statistics and analysis

The data was analyzed with estimation methods to calculate mean, mean differences, confidence intervals ^93^, and Hedges’ *g* where appropriate ^62^. Data are presented for each individual fly as well as the mean difference in estimation plots. 95% confidence intervals for the mean differences were calculated using bootstrap methods (resampled 5000 times, bias-corrected, and accelerated) and are displayed with the bootstrap distribution of the mean. Effect size was measured using Hedges’ g and are as per standard practice referred to as either ‘trivial’ (*g* < 0.2), ‘small’ (0.2 < *g* < 0.5), ‘moderate’ (0.5 < *g* < 0.8) or ‘large’ (*g* > 0.8)^94^. Hedges’ is a quantitative measurement for the difference between means, and is an indication of how much two groups differ from each other where a Hedges’ g of 1 shows that the two groups differ by 1 standard deviation. Commonly used significance testing and power calculations were avoided following recommended practice ^94,95^ the Mann Whitney *U* statistic was used to calculate *P* values for *pro forma* reporting exclusively. To indicate estimate precision, 95% confidence intervals (95CI) were calculated using bootstrap methods and reported in text and/or as error bars ^94^. The behavioral data are presented as mean-difference estimation plots. Confidence intervals were bias-corrected and accelerated, and are displayed with the bootstrap distribution of the mean; resampling was performed 5,000 times. While P values reported are related to two-tailed Student’s t-test statistics, significance testing was not performed ^93^.

## Author Contributions

*Conceptualization*: ACC, MH; *Methodology*: AE, JG, PA; JCS (activity assay development) *Investigation*: AE (thermosensation, feeding assay, activity, climbing, iTASSER modelling), JG (transcriptomics and toxin identification); PA, CG, EH: (synthesis, oxidative folding and purification of peptides Tv1 and Tsu1.1); *Data Analysis*: AE, PA, JG, MH, ACC, JCS (conotoxin comparison, thermosensation, feeding assay, activity, climbing); *Writing - Original Draft*: AE; *Writing - Revision*: AE, PA, JG, MH and ACC; *Visualization*: AE, PA, JG, MH and ACC; *Supervision*: MH and ACC; *Project Administration*: ACC and MH; *Funding Acquisition*: MH and ACC.

## Acknowledgements

We thank Dr. Joses Ho for the scripts to calculate and plot the data. We thank Si Yi Lim for providing help with pain data acquisition and food intake analysis.

